# Genomic signatures of selection associated with benzimidazole drug treatments in *Haemonchus contortus* field populations

**DOI:** 10.1101/2022.04.05.487096

**Authors:** Janneke Wit, Matthew L. Workentine, Elizabeth Redman, Roz Laing, Lewis Stevens, James A. Cotton, Umer Chaudhry, Qasim Ali, Erik C. Andersen, Samuel Yeaman, James D. Wasmuth, John S. Gilleard

**Affiliations:** Department of Comparative Biology and Experimental Medicine, Host -Parasite Interactions (HPI) program, University of Calgary, Calgary, Alberta, Canada; Faculty of Veterinary Medicine, University of Calgary, Calgary, Alberta, Canada; Institute of Biodiversity Animal Health and Comparative Medicine, College of Medical, Veterinary and Life Sciences, University of Glasgow, Garscube Campus, Glasgow, UK; Tree of Life, Wellcome Sanger Institute, Cambridge, UK; Wellcome Sanger Institute, Wellcome Genome Campus, Hinxton, Cambridgeshire, UK. CB10 1SA.; University of Edinburgh, Roslin Institute, Easter Bush Veterinary Centre, Roslin, Midlothian, UK; Department of Veterinary Parasitology, University of Veterinary and Animal Sciences Lahore, Pakistan; Molecular Biosciences, Northwestern University, Evanston, IL, USA; Department of Biological Sciences, University of Calgary, Calgary, Alberta, Canada; Department of Ecosystem and Public Health, Faculty of Veterinary Medicine, University of Calgary, Calgary, Alberta, Canada

## Abstract

Genome-wide methods offer a powerful approach to detect signatures of drug selection in parasite populations in the field. However, their application to parasitic nematodes has been limited because of both a lack of suitable reference genomes and the difficulty of obtaining field populations with sufficiently well-defined drug selection histories. Consequently, there is little information on the genomic signatures of drug selection for parasitic nematodes in the field and on how best to detect them. This study was designed to address these knowledge gaps using field populations of *Haemonchus contortus* with well-defined and contrasting benzimidazole-selection histories, leveraging a recently completed chromosomal-scale reference genome assembly. We generated a panel of 49,393 ddRADseq markers and used this resource to genotype 20 individual *H. contortus* adult worms from each of four *H. contortus* populations: two from closed sheep flocks that had an approximately 20-year history of frequent treatment exclusively with benzimidazole drugs, and two populations with a history of little or no drug treatment. The populations were chosen from the same geographical region to limit population structure in order to maximize the sensitivity of the approach. A clear signature of selection was detected on the left arm of chromosome I centered on the isotype-1 β-tubulin gene in the benzimidazole-selected but not the unselected populations. Two additional, but weaker, signatures of selection were detected; one near the middle of chromosome I and one near the isotype-2 β-tubulin locus on chromosome II. We examined genetic differentiation between populations, and nucleotide diversity and linkage disequilibrium within populations to define these two additional regions as encompassing five genes and a single gene. We also compared the relative power of using pooled versus individual worm sequence data to detect genomic selection signatures and how sensitivity is impacted by sequencing depth, worm number, and population structure.

In summary, this study used *H. contortus* field populations with well-defined drug selection histories to provide the first direct genome-wide evidence for any parasitic nematode that the isotype-1 β-tubulin gene is the quantitatively most important benzimidazole resistance locus. It also identified two additional genomic regions that likely contain benzimidazole-resistance loci of secondary importance. Finally, this study provides an experimental framework to maximize the power of genome-wide approaches to detect signatures of selection driven by anthelmintic drug treatments in field populations of parasitic nematodes.

**AUTHOR SUMMARY:** Benzimidazoles are important anthelmintic drugs for human and animal parasitic nematode control with ∼0.5 billion children at risk of infection treated annually worldwide. Drug resistance is common in livestock parasites and a growing concern in humans. *Haemonchus contortus* is the most important model parasite system used to study anthelmintic resistance and a significant livestock pathogen. It is also one of the few parasitic nematodes with a chromosomal-scale genome assembly. We have undertaken genome-wide scans using a dense RADseq marker panel on worms from natural field populations under differing levels of benzimidazole selection. We show that there is a single predominant genomic signature of selection in *H. contortus* associated with benzimidazole selection centred on the isotype-1 β-tubulin locus. We also identify two weaker signatures of selection indicative of secondary drug resistance loci. Additionally, we assess the minimum data requirements for parameters including worm number, sequence depth, marker density needed to detect the signatures of selection and compare individual to Poolseq analysis. This work is the first genome-wide study in a parasitic nematode to provide direct evidence of the isotype-1 β-tubulin locus being the single predominant benzimidazole resistance locus and provides an experimental framework for future population genomic studies on anthelmintic resistance.

## 1. INTRODUCTION

Parasitic nematode infections are of major medical and agricultural importance worldwide [1,2]. Control strategies largely depend on the regular use of anthelmintic drugs, which has caused widespread anthelmintic drug resistance in many parasitic nematode species of livestock [1,3]. Additionally, concerns are increasing that mass drug administration programs are selecting for anthelmintic resistance in human helminths [4–8]. Over the last 30 years, there has been a large amount of research into the molecular genetic basis of anthelmintic resistance, particularly in gastrointestinal nematodes of livestock [9]. Most of this work has been dominated by candidate gene studies, where certain genes are prioritized based on knowledge of the drug mode-of-action [10–14] or by the extrapolation of genetic studies in the model organism *Caenorhabditis elegans* [15,16]. Whilst this approach has been successful in some cases, most notably for the benzimidazole drug class, it has been much less successful in others [17]. For example, in the case of the macrocyclic lactones, which is one of the most important broad spectrum classes of anthelmintic drugs, the evidence implicating the leading candidate genes has been inconsistent between studies [18–22].

Genome-wide approaches, which make no *a priori* assumptions about the underlying mechanisms of resistance, are potentially more powerful approaches to identify anthelmintic resistance loci than candidate gene studies. The most successful example in a helminth to date was the use of classical linkage mapping to identify a Quantitative Trait Locus (QTL) for oxamniquine (OXA) resistance in the human trematode *Schistosoma mansoni* [23]. Subsequent work confirmed the functional importance of a mutation in a sulfotransferase enzyme (SmSULT-OR) gene to the OXA-resistant phenotype [23,24].

The small ruminant parasite *Haemonchus contortus* is the leading parasitic nematode model for anthelmintic resistance research for a variety of reasons, including the availability of well characterized anthelmintic resistant isolates, a good understanding of its genetics, and the ability to undertake genetic crosses [9,25–27]. A chromosomal-scale reference genome assembly and annotation enables genome-wide approaches in this parasitic nematode species (Doyle et al 2020). This new reference genome was successfully used to map the major ivermectin resistance QTL on chromosome V in two independent laboratory passaged *H. contortus* strains by a serial backcrossing approach [28,29]. However, such genetic crosses are extremely challenging in parasitic nematodes compared to model organisms and have only been demonstrated in a few parasite species [26,30]. An alternative way to map the genetic loci underlying anthelmintic resistance is to use population genomic approaches on field populations. This approach is not only technically simpler and more scalable but is also more likely to identify anthelmintic resistance loci that are relevant to field populations.

As reference genomes for parasitic nematodes continue to improve, and DNA sequencing costs reduce, population genomic studies on natural field populations should become increasingly feasible. However, there is currently a lack of data from field populations with sufficiently well-defined and contrasting drug treatment histories to unambiguously link signals of selection in the genome with specific drug treatments. For example, most *H. contortus* field populations have been subject to treatment with multiple different drug classes and to significant animal movement. Consequently, there is a need for studies on field populations with well-defined drug treatment histories to elucidate the genomic location and nature of signatures of selection associated with specific drug classes.

It is now well established that mutations in the isotype-1 β-tubulin gene are important determinants of benzimidazole resistance in *H. contortus* and other strongylid nematodes [10,31–36]. However, a key knowledge gap at present is whether the isotype-1 β-tubulin locus is the only major locus under selection by benzimidazole drug treatments or whether there are additional selected loci and, if so, their relative importance [37]. Interestingly, a recent study in which whole-genome sequencing was applied to 233 individual adult *H. contortus* worms from field populations, with varied and complex drug selection histories, identified multiple regions of the genome that showed signatures of selection including around the isotype-1 β-tubulin locus [38]. However, this study could not associate these signatures of selection with treatment by a specific drug class as the populations used had a variety of complex drug treatment histories. A major aim of our current study was to determine which loci in the *H. contortus* genome show strong evidence of selection that are specifically associated with benzimidazole selection in the field.

Another aim of this study was to take advantage of the clear and contrasting benzimidazole drug selection histories of the *H. contortus* populations on government and rural farms in Pakistan to investigate the best technical approach to detect genomic signatures of drug selection in field populations. Reduced-representation sequencing methods, where a random but consistent subset of the genome is sequenced, are potentially powerful and cost-effective for genome-wide methods [39]. Larger numbers of organisms and/or samples can be sequenced than by whole-genome sequencing at the same cost, and while sequencing costs are dropping, cost is an important consideration when examining larger sample sets, particularly for laboratories outside of the major genome sequencing centres [40,41]. One of the more widely used reduced-representation approaches is Restriction-site Associated DNA sequencing (RADseq) [42–44]. This approach has been extremely successful at investigating population diversity and differentiation in a wide range of organisms [45–48] but it is still a challenge to get a high-density marker set to identify signatures of selection [49–51].

Here, we describe the development of a high density RADseq marker-set, and its use to investigate the genomic signature(s) of benzimidazole drug selection in *H. contortus* field populations. The strongest signature of benzimidazole drug selection, by far, that was present in two populations with a history of regular benzimidazole treatments, but absent in two untreated populations, was in the genomic region surrounding the isotype-1 β-tubulin gene. Two other genomic regions under drug selection were identified but the signals of selection at these loci were weaker. In addition, we investigated the effect of sample number and read depth, and compared the analysis of individual worm sequence data with pooled sequence data on the ability of RADseq to detect genomic selection signatures in order to provide a framework for future experimental design.

## 2. MATERIALS AND METHODS

### 2.1 Parasite material

Populations of adult *H. contortus* worms were harvested from four different sheep flocks in the Punjab province of Pakistan [52]. Two populations were sampled from government farm flocks with a history of regular BZ treatment over several decades; Treated 1 (T1) from Okara and Treated 2 (T2) from Jahangirabad (T1 is Pop1S and T2 is Pop3S in [52]). These government farms established their sheep flocks using local sheep breeds (lohi and kajli) in 1985 and 1989 respectively and have been alternatively treated with albendazole and oxfendazole, but not other anthelmintic classes, approximately every three months since their establishment. These herds have been closed to animal movement since they were established. There has been some historical movement of stock between the farms, but no animals have been introduced from elsewhere. Two other populations were sampled from rural flocks with no history of drug treatment, from two abattoirs in the same region; Untreated 1 (U1) and Untreated 2 (U2). Worms were harvested directly from ewe abomasa at necropsy at abattoirs from sheep carcases following routine slaughter for human consumption and stored in 70% ethanol at -80°C. For this study, 20 females were randomly selected from each sample.

### 2.2 DNA extraction

Adult female worms were rehydrated in sterile water for 60 minutes prior to extraction. Tissue was isolated from the anterior portion of the adult worm, avoiding the gonads containing progeny, and DNA was extracted using a protocol adapted from Bennet *et al.* [53]. After dissection of the animal, the anterior portion was transferred to 600 µl lysis buffer (384 µl H2O, 60 µl 1M Tris HCl (pH 8.5), 60 µl 1% SDS, 60 µL 0.5M EDTA (pH 7.5), 12 µl 5M NaCl, 12 µl proteinase K (14-22 mg/mL, Qiagen), and 12 µl β– mercaptoethanol). Samples were incubated overnight at 55°C before adding 2 µl of RNaseA (100 mg/ml, Qiagen) and incubated for 15 min at 45°C. After adding 150 µl 5M NaCl and 100 µl CTAB/NaCl samples were incubated at 65°C for 15 min. An equal volume of phenol:chloroform:isoamyl alcohol (25:24:1, Sigma) was added and the solution was incubated for 1.5 hr on a slow rocker at room temperature (RT). Samples were centrifuged at 3000 g for 15 min at 12°C and the aqueous phase was collected. An equal volume of chloroform:isoamyl alcohol (24:1, Sigma) was added to the aqueous phase and the solution incubated for 1 hour at RT, centrifuged as before and the aqueous phase collected. Two volumes of ice-cold ethanol were added and gently mixed. The solution was placed at -20°C overnight before being centrifuged as before at 4°C. After removal of the supernatant, the pellet was washed twice with 70% EtOH. After the final wash, the sample was centrifuged at 14000 rpm for 10 min, the supernatant removed, centrifuged for an additional minute to remove the remainder of the ethanol and the resulting pellet air dried before resuspension in 30 µl water and incubation overnight at 4°C. Sample quality was checked by quantification on the Qubit system (ThermoFisher) and A_260_/A_280_ absorbance ratios on a NanoVue (Biochrom Spectrophotometers). 2.5 µl per sample (46.4-400 ng) was added to a REPLI-g Single Cell Kit (Qiagen) for whole genome amplification (WGA) following the manufacturers protocol for Amplification of Purified Genomic DNA.

### 2.3 Amplicon sequencing of isotype-1 β-tubulin and isotype-2 β-tubulin genes and haplotype network analysis

Amplicon sequencing was used to determine the frequency of single nucleotide polymorphisms (SNPs) in the isotype-1 and isotype-2 β-tubulin genes previously associated with benzimidazole resistance in *H. contortus* - F200Y (TTC>T**<u>A</u>**C), F167Y (TTC>T**A**C), E198A (GAA>G**C**A), or E198L (GAA>**TT**A) – in each population. The amplicon sequencing protocol and data analysis has been previously described in Avramenko et al. [54,55] and adapted for isotype-2 -tubulin with primers in **File S1**. For the haplotype analysis of isotype-1 β-tubulin, reads classified as *H. contortus* by the Mothur [56] pipeline in the amplicon data analysis were extracted from the original FASTQ files. Primers were removed with Cutadapt [57], and haplotype analysis was conducted with DADA2 [58,59], following their pipeline tutorial for demultiplexed Illumina paired-end reads. Resulting amplicon sequence variants (ASVs) were used to create a haplotype network using ape and pegas in R [60,61].

### 2.4 Preparation and sequencing of double-digest single worm RAD libraries

The single worm WGA samples were prepared for sequencing following a protocol adapted from Peterson *et al.* (2012). Post-WGA samples were purified using a standard bead clean-up protocol using AMPure XP beads (Beckman), with 1.5x beads instead of 1.8x sample:bead volume, and eluted in water. The DNA was digested with *MluC*I (5µl Cut smart buffer, 1.5-2 µg DNA, 3 µl *MluC*I (New England Biolabs Inc. (NEB), and water to make up 49 µl) for three hours at 37°C after which 3 µl *Nla*II (NEB) was added and the reaction incubated at 37°C for a further three hours. The samples were cleaned as before. Barcoded adapters were ligated in a 50 µl reaction volume, with 4 µl T4 buffer, 7.5 µl each of adapter P1 and P2, 0.7 µl T4 ligase (2 M U/ml, NEB), 250 ng DNA (adapters in **File S1**) and water. Libraries from individual worms of different populations were randomized to create pools containing 20 individuals each for sequencing, cleaned as before, and size-selected on a PippinPrep (Sage Science) using the standard protocol for Tight Range (190 bp) on a 3% internal standards cartridge, resulting in a fragment range of 190-300 bp with a peak at 210 bp. Quality and fragment size was checked on a Tape Station (Agilent), and P5 and P7 adapters were ligated per pool following the KAPA HiFi Hotstart PCR kit standard protocol (Kapa Biosystems). Pooled samples were then combined into one sample and cleaned as before. Because the fragments ranged in size, quantity was checked in triplicate using a Qubit broad range kit, rather than a qPCR approach, and diluted to 4 nM after correction for average size.

The pooled individual worm libraries were sequenced on the Illumina NextSeq platform. Populations U1, U2, T1, and T2 were sequenced on one High Output 2 x 75 bp kit. The U2 population was discarded after only 20.22 - 53.54 % of its reads were mapped to the reference with the remainder being bacterial contamination. Consequently, the U2 population was re-sequenced using a Mid Output 2 x 75 bp kit. The loading concentration was 1.2 pM, and 50% PhiX was spiked in for diversity due to the low diversity library.

Fragment-size selection was based on predicted fragment recovery with the draft genome assembly (GCA_000469685.1 [62]) using SimRAD [63].

### 2.5 Quality Control and Analysis of Sequence Data

The data were demultiplexed using process_radtags in the Stacks pipeline [64] (SRA accession number PRJN822658). All samples were assessed for general sequence quality using FastQC [65], before removing adapters using Cutadapt [57]. Read quality was high across the full read length for both sequencing runs; therefore, no hard trimming was conducted. Reads were aligned against the chromosomal-scale *H. contortus* reference genome assembly (GCA_000469685.2, N50 = 47.38 Mb, N50(n) = 3, n= 7 [66]) using Bowtie2 (--local --very-sensitive-local -X 300 -I 80 [67]). A BAM file was generated for each sample using Samtools by filtering for paired reads only and then sorting [68]. The sorted data was analyzed using gstacks and populations using the –fstats flag from the Stacks pipeline. Only SNPs that were present in three of four populations, in 50% of individuals per population, and at a minimum frequency of 5% were retained in the output. In addition, only the first SNP per RAD locus was reported to avoid linkage and variable SNP count within a locus to affect the interpretation of the data.

To assess the robustness of the RADseq analysis and investigate the limits to its ability to detect signatures of selection, additional analyses were conducted using the same basic parameters but fewer individual worms per population (5, 10, and 15, randomly selected), as well as fewer reads per individual (1M, 1.5M, and 2M reads where possible, randomly selected using seqtk sample). Finally, the raw reads were realigned to an earlier version of the reference genome (GCA_000469685.1, N50 = 0.099 Mb, N50(n) = 876, n= 12,915 [62]), to illustrate the effect of genome quality on identifying regions of interest in the genome.

### 2.6 Population differentiation, expected heterozygosity, and linkage disequilibrium analyses to detect signatures of selection

All analyses were performed on data generated by Stacks (version 2.2), reporting one SNP per RAD-locus, unless otherwise stated. To examine overall population differentiation, population level F_ST_ values were calculated and a Principal Component Analysis was conducted on all reported SNPs across the populations using the vcfR (version 1.8.0) and adegenet (version 2.1.1) packages in R [69,70].

To detect regions in the genome where population differentiation was significantly higher than background noise, we used a top candidate method as described in Yeaman et al. [71]. The 99^th^ percentile of all F_ST_ values was determined and F_ST_ values were then binned over 150 kb intervals. Top candidate regions were identified as bins with an exceptional proportion of their total SNPs being F_ST_ outliers based on a binomial distribution. For expected heterozygosity, a similar approach was undertaken. Outliers were defined as expected heterozygosity values falling in the lowest 25^th^ percentile and then binned across 150 kb regions. Top candidate bins were identified as those having a higher than expected proportion of low expected heterozygosity based on a binomial distribution. Linkage disequilibrium per population was calculated in R using the cor() function with “pairwise.complete.obs”. These values were binned across 150 kb regions and the percentage of R^2^ values above the 99^th^ percentile of all R^2^’s per population was recorded.

### 2.7 Analysis of mock pooled sequence data

All RADseq data from individual worms per population were pooled to create mock pooled sequence data. Data were pooled after demultiplexing with process_radtags, as these RAD-specific barcodes would not be present in Illumina sequence data of pooled samples. Samples were then aligned with Bowtie2 as above. The resulting aligned files were subsampled randomly using seqtk, resulting in 100⨉ genome coverage. For analyses of individual samples, average expected heterozygosity was determined for one SNP per RAD-locus, therefore bedtools was used to concatenate SNPs within 150 bp of each other to one marker for pooled samples [72,73]. These subsamples were analyzed using PoPoolation Variance-sliding.pl (Variance-sliding.pl --measure pi --min-count 2 -- min-qual 20 --min-coverage 15 --max-coverage 400 --pool-size 20 --window-size 1 -- step-size 1) and Popoolation2 snp-frequency-diff.pl (snp-frequency-diff.pl --min-count 3--min-coverage 15 --max-coverage 400 [74,75], generating Tajima’s Pi and F_ST_ statistics.

### 2.8 Detecting genes in genomic regions under selection

All genes and corresponding proteins within the genomic regions showing consistent signatures of selection in the treated populations across multiple estimates were extracted from the gene annotation file haemonchus_contortus.PRJEB506.WBPS14.annotations.gff3 and the FASTA file haemonchus_contortus.PRJEB506.WBPS14.protein.fa. Potential causal variation in these genes would not necessarily be identified in RAD-data as a result of limited sequence coverage of the region. Therefore functional annotation of the selected genes was performed with InterPro version 5.35-74.0 and Pfam database version 32.0 (interproscan.sh -i [input_protein.fa] -d out -dp -t p --goterms -appl Pfam -f tsv).

### 2.9 Data availability

All samples for the ddRADseq project are collected under the BioProject accession number PRJNA822658.

## 3. RESULTS

### 3.1 Developing a panel of genome-wide RADseq markers for *H. contortus*

Genome-wide RADseq markers were generated by digestion of the genome with *MluC*I and *Nla*II, fragment size selection of 190-300 bp (peak at 210 bp), and sequencing on an Illumina NextSeq (2x75bp). On average, 77.7% of the reads mapped to the reference genome sequence (max: 81.4%, min: 60.2%, median: 78.6%). Of the mapped reads, 77.6% passed the paired-read filter providing an overall average total of 60.33% of raw reads for further analysis using Stacks [64] number of reads kept: average = 3,337,301, median = 3,186,605, min = 1,283,698, max = 5,501,942). We kept only those markers present in 50% of individuals per population and in all three populations (75.4% of all primary alignments). Average coverage and marker density for all comparisons are shown in **Table 1** and **Fig. S1**. With baseline parameters, the range of reads per individual was 4.4X-14.2X coverage.

**Table 1.** Overview of coverage and average marker recovery across different analysis parameters and data subsets.

| Dataset | Average coverage<br>per marker | Marker count |
| --- | --- | --- |
| ***Mapping to reference genome:*** |  |  |
| ***GCA_000469685.2*** |  |  |
| Baseline (50% individuals, 3 populations) | 9.1 | 49,393 |
| Stringent (75% individuals, 3 populations) | 9.1 | 26,269 |
| ***Mapping to reference genome:*** |  |  |
| ***GCA_000469685.1*** |  |  |
| Baseline (50% individuals, 3 populations) | 9.8 | 42,987 |
| ***Data Subsets (50% individuals, 3 populations)*** |  |  |
| Population size = 15 | 9.0 | 45,226 |
| Population size = 10 | 9.4 | 49,687 |
| Population size = 5 | 9.0 | 52,532 |
| Read depth = 2.0M/sample | 7.4 | 42,818 |
| Read depth = 1.5M/sample | 6.3 | 37,432 |
| Read depth = 1.0M/sample | 5.0 | 30,136 |
| Read depth = 0.5M/sample | 3.3 | 18,283 |
| Read depth = 0.25M/sample | 2.3 | 7,769 |
| Read depth = 0.1M/sample | 1.6 | 1,666 |

### 3.2 Population-wide genetic diversity and isotype-1 β-tubulin resistance allele frequencies

Genome-wide pairwise F_ST_ was low between all populations, based on population differentiation across 43,587-48,227 markers (Table 2). Population differentiation was highest in the pairwise comparisons involving population U2 (**Table 2**). PCA illustrated the genetic similarity of the two drug-treated populations (T1 and T2), with some overlap with U1 but not U2, consistent with the F_ST_ results (**Fig. 1B**). Removing variation on chromosome I (which contains the isotype-1 β-tubulin gene) decreases differentiation between U1 and T1 and T2 (**Fig. 1C**). Although U2 was collected in the Punjab province, it was collected in Sargodha, about 200 km from Lahore, where the other populations were collected. This distance potentially explains the slightly increased levels of differentiation between U2 and the other populations (**Fig. 1A**).

**Fig. 1.**
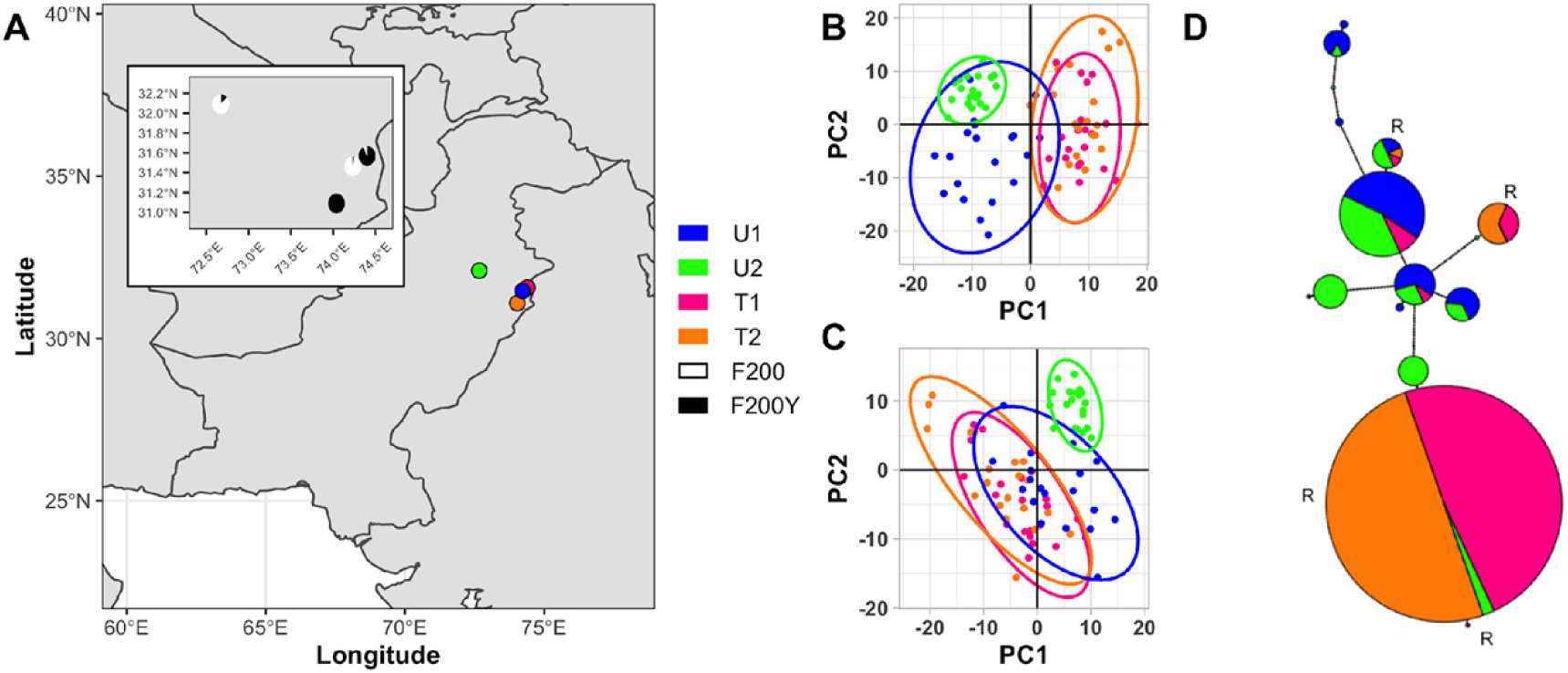
**A)** Sampling location, inset shows the percentage of P200Y mutations of the isotype-1 β-tubulin locus. **B)** PCA plot comparing SNP data of the four populations. **C)** PCA plot comparing SNP data of the four populations without chromosome I. **D)** Haplotype network of amplified sequence variants (ASVs) of the isotype-1 beta-tubulin locus amplicon data.

**Table 2.** Pairwise F_ST_ values between populations.

| Population | T1 | T2 | U1 | U2 |
| --- | --- | --- | --- | --- |
| T1 |  | 0.023 | 0.031 | 0.047 |
| T2 |  |  | 0.033 | 0.046 |
| U1 |  |  |  | 0.037 |

Of the isotype-1 β-tubulin SNPs previously associated with BZ resistance in *H. contortus,* only 200Y (TTC > T**<u>A</u>**C) was detected in the populations after genotyping by amplicon sequencing. The frequencies of F200Y in the U1, U2, T1, and T2 populations were 2.5%, 12.5%, 100%, and 92.5%, respectively, consistent with their drug treatment history and indicative of their benzimidazole-resistance status (**Fig. 1A inset**). The most common resistance isotype-1 β-tubulin haplotypes are present at high frequency in both T1 and T2 populations (**Fig. 1D**). The isotype-2 β-tubulin gene was also genotyped by amplicon sequencing for all individual worms to identify potential candidate resistance mutations at codons 167, 198, and 200. All worms were homozygous for “susceptible alleles” (198E, and 200F), except for one worm in population U1 which was heterozygous for a F200Y (TTC > T**<u>A</u>**C) allele.

### 3.3 The major region of genetic differentiation between treated and non-treated populations centers around the isotype-1 β-tubulin locus on chromosome I

All four pairwise genome-wide comparisons between a treated and an untreated population showed a major region of elevated F_ST_ on chromosome I, centered around the isotype-1 β-tubulin locus (chromosome I, around 7.0Mb, **Fig. 2A**). The size of this region depended on the particular pairwise comparison but was between 3.5 and 10 Mb.

**Fig. 2.**
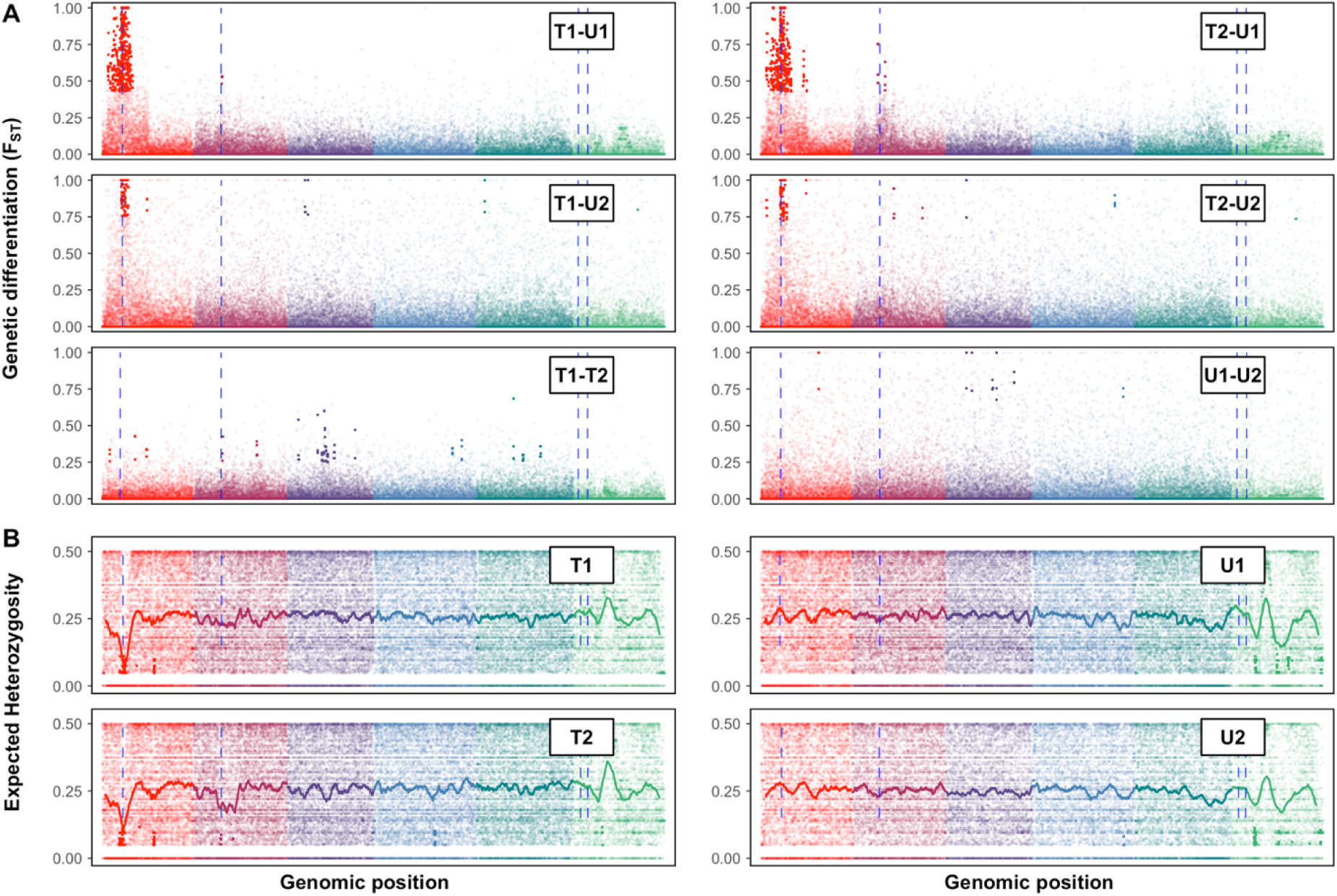
**A)** Genome-wide pairwise F_ST_ **B)** Genome-wide expected heterozygosity estimates per population. Chromosomes are indicated by different colors and ordered from I to V with X on the far right in green. Bolded data points represent loci with an F_ST_ significantly different from background noise, as described by the top candidate test [71] and the Materials and Methods. The blue lines indicate the location of isotype-1 β-tubulin on chromosome I, isotype-2 β-tubulin on chromosome II, and isotype-3 and 4 β-tubulin on chromosome X. The solid line in B is the rolling average across 500 RAD loci.

Pairwise comparisons between either the two treated or two untreated populations did not have an elevated F_ST_ in this region (**Fig. 2A**). In addition, a weaker region of elevated F_ST_ was centred on the isotype-2 β-tubulin locus for the T2-U1 and T2-U2 but not the T1-U1 and T1-U2 pairwise comparisons.

### 3.4 Patterns of expected heterozygosity and linkage disequilibrium indicate a strong signature of selection around the isotype-1 β-tubulin locus and identify a second locus under selection on chromosome I

In the treated but not the untreated populations, the region surrounding the isotype-1 β-tubulin gene, which is located at 7,027,095 Mb on chromosome I, has distinctly reduced expected heterozygosity (position range 5,708,200 Mb - 8,113,930 Mb) and elevated linkage disequilibrium (position range 6,075,000 Mb – 8,325,000 Mb) (**Fig. 2B and Fig. 3**). There is a second region on chromosome 1 with a distinct decrease in expected heterozygosity (position range 26,568,594 Mb - 26,676,298 Mb) and an elevated linkage disequilibrium (position range 26,025,000 Mb - 26,625,000 Mb) (**Fig. 3**). A third region on chromosome 2 has a region of reduced expected heterozygosity in treated population T2 (position range 13,221,871 Mb – 13,270,135 Mb) close to the isotype-2 β-tubulin gene (which is located at 13,433,296 Mb) (**Fig. 2B**). Linkage disequilibrium around the third region is not elevated (**Fig. S2**).

**Fig. 3.**
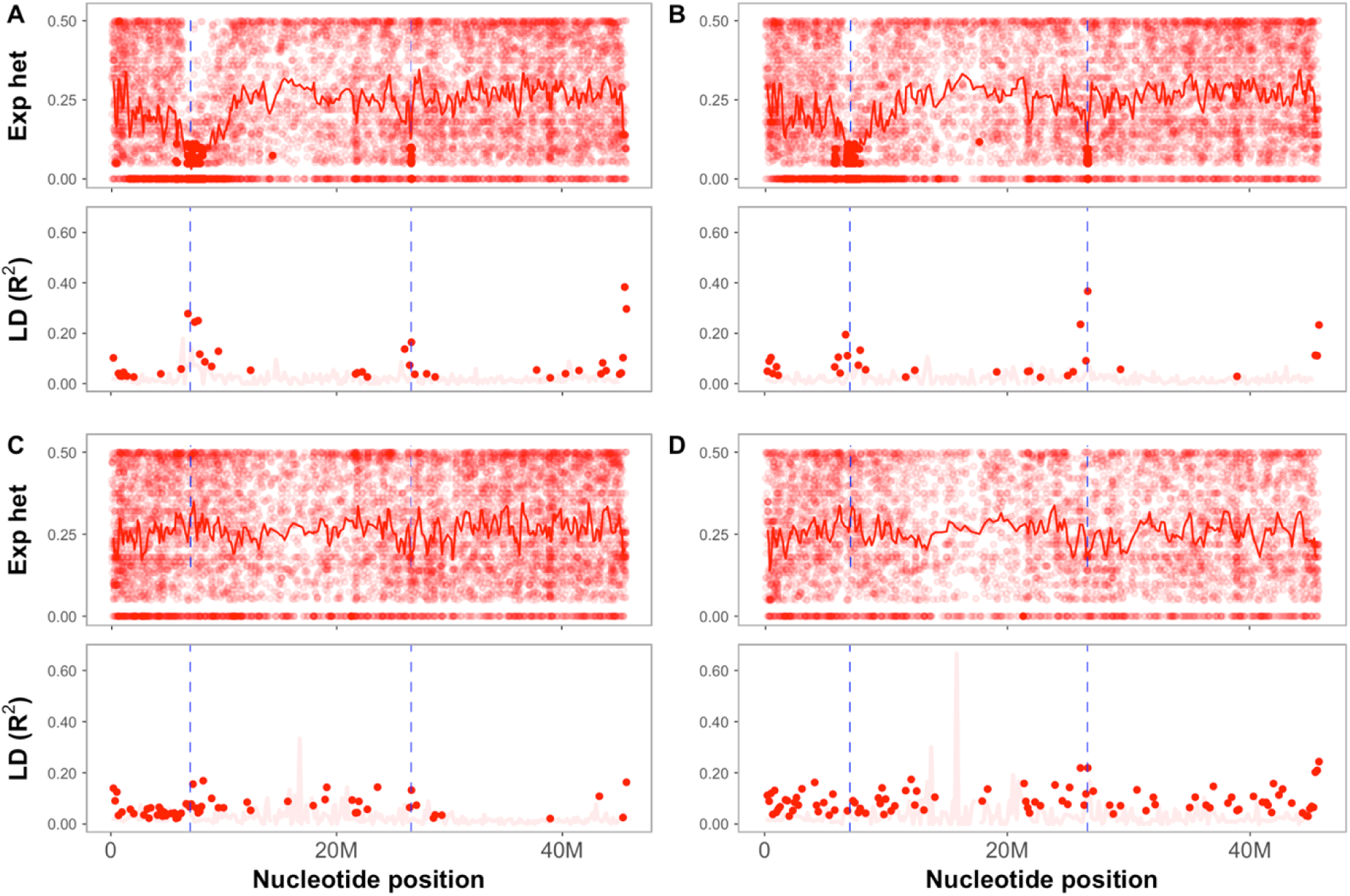
Chromosome I expected heterozygosity (Exp het), and linkage disequilibrium (R^2^ **A)** Population T1 **B)** Population T2 **C)** Population U1, and **D)** Population U2. Bolded data points represent loci significantly different from background noise, as described by the top candidate test [71] and the Materials and Methods. Opaque points and lines indicate non-significant loci. The two dashed lines indicate isotype-1 β-tubulin (left, around 7027095 Mb) and the second locus on chromosome I (right, around 26596797 Mb). The solid line in expected heterozygosity plots is the rolling average across 500 RAD-loci.

### 3.5 Effect of sequencing depth, sample size, population structure, and genome assembly quality on the signature of selection detected by F_ST_

We investigated the amount of RADseq data required to detect the signature of selection, as measured by the elevation in F_ST_ by randomly sub-sampling the dataset. As expected, decreasing the numbers of individuals per population in the analysis decreases the signal of elevated F_ST_ around the isotype-1 β-tubulin gene. However, the signal remains detectable even when data from as few as five individual worms are included in the analysis, especially for the comparisons between a treated population (T1 or T2) and the U1 untreated population (**Fig. 4A**, **Fig. S3**). Decreasing the number of reads per individual only minimally affected the elevation of F_ST_ around the isotype-1 β-tubulin locus until the coverage drops below 3.3X (read depth 0.5M), particularly in comparisons with the more genetically divergent U2 population (**Table 1, Fig. 4B**, **Fig. S4**).

**Fig. 4.**
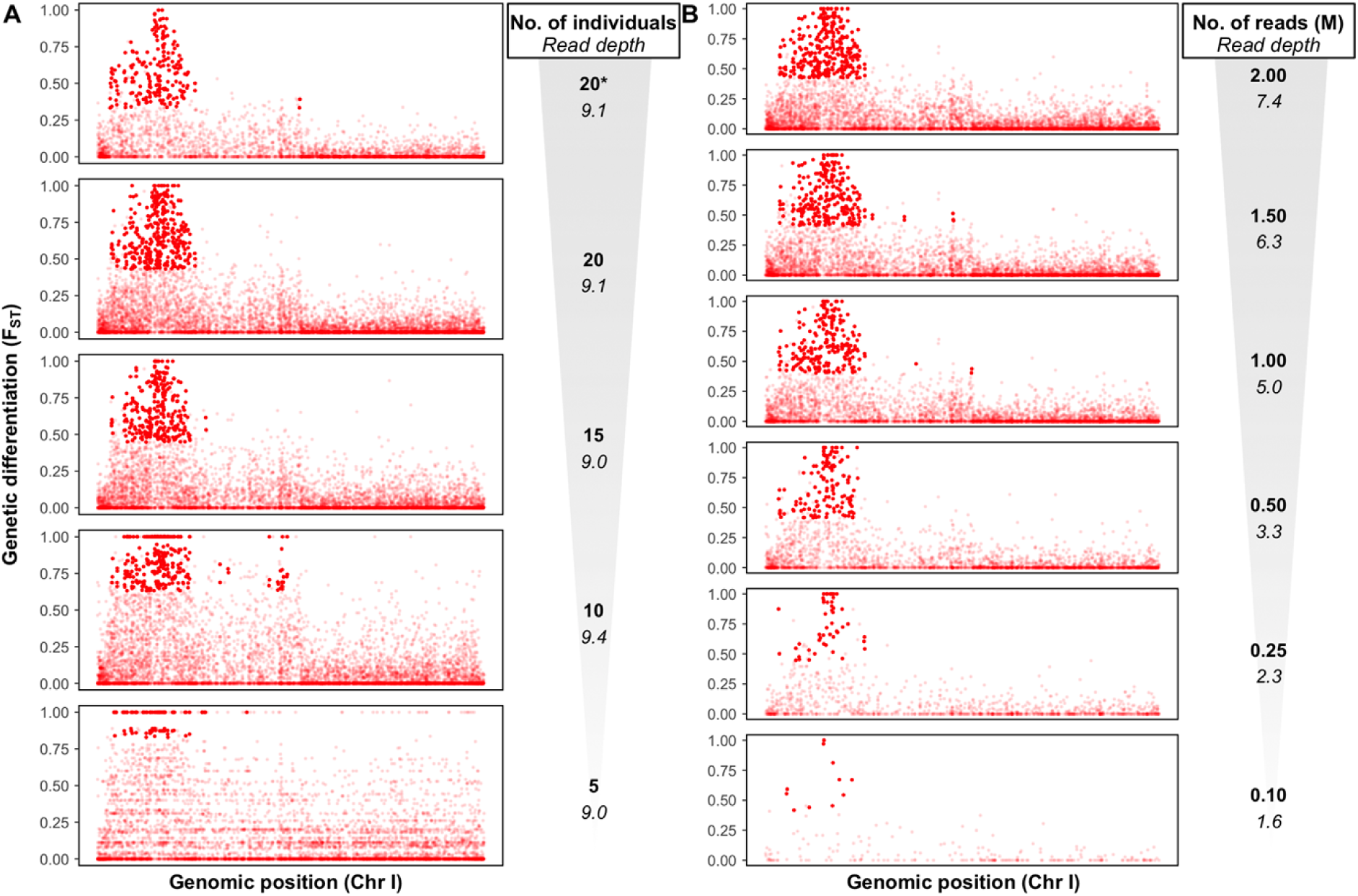
F_ST_ analysis between T1 and U1. Each dot represents a variable locus between the paired populations, solid colors represent loci with an F_ST_ significantly above background noise, as described by the top candidate method [71] **A)** F_ST_ estimates across chromosome I with decreasing numbers of individuals per population. 20*; stringent, variants are called when represented in 75% of individuals per population. 20, 15, 10 and 5; variant data present for 50% of the individuals per population, 20-15-10-5 **B)** F_ST_ estimates across chromosome I for decreasing numbers of reads per individual sample. Variant data had to be present in 50% of the individuals per population, but only the indicated number of reads was used per sample. Read depth indicated under read count.

### 3.6 Gene models within the regions defined by the signals of selection in populations T1 and T2

With the chromosomal-scale reference genome, the RADseq data was used to detect genomic regions with signatures of selection shared by the treated populations. With the gene annotations that accompany this genome, we further explored these regions for genes related to BZ resistance. We selected three regions of interest based on the results of the four estimates. The narrowest ranges for these three regions, based on an intersection of the detected ranges in all comparisons are (1) chromosome I: 6,450,000-8,113,930 Mb; (2) chromosome I: 26,568,594-26,625,000 Mb; and (3) chromosome II: 13,221,871-13,270,135 Mb. In total, these regions contain 161, five, and one gene(s), respectively (**S2 File**). As discussed, region 1 contains isotype-1 β-tubulin (HCON_00005260). Region 2 contains five genes; HCON_00018620, HCON_00018630, HCON_00018640, HCON_00018650, and HCON_00018660. Region 3, which is based on diversity estimates only, contains a single gene (HCON_00043570) which is predicted to be a hyaluronidase.

### 3.7 Ability to detect signatures of selection using pooled sequence analysis

We compared the ability of a pool-seq approach to detect the signature of selection to that of the single worm approach by pooling the individual worm RADseq data for each population. Using Popoolation2, 32,234 SNPs were identified as differential in at least one pairwise population comparison (**Fig. 5C**, **Fig. S5,** [74]). These pairwise comparisons show a peak in F_ST_ around the isotype-1 β-tubulin gene, especially for comparisons between both treated populations and U1, with a less clear peak in the comparison with U2. Diversity (π) was determined with all available SNP data, with 27,719-113,917 estimates across the genome. These loci were then analysed with Popoolation using the loci as a gene map, with 5,764-14,156 π estimates per population. None of the regions under selection as detected with individual samples were identified with diversity estimates in the pooled analysis (**Fig. S6**).

**Fig. 5.**
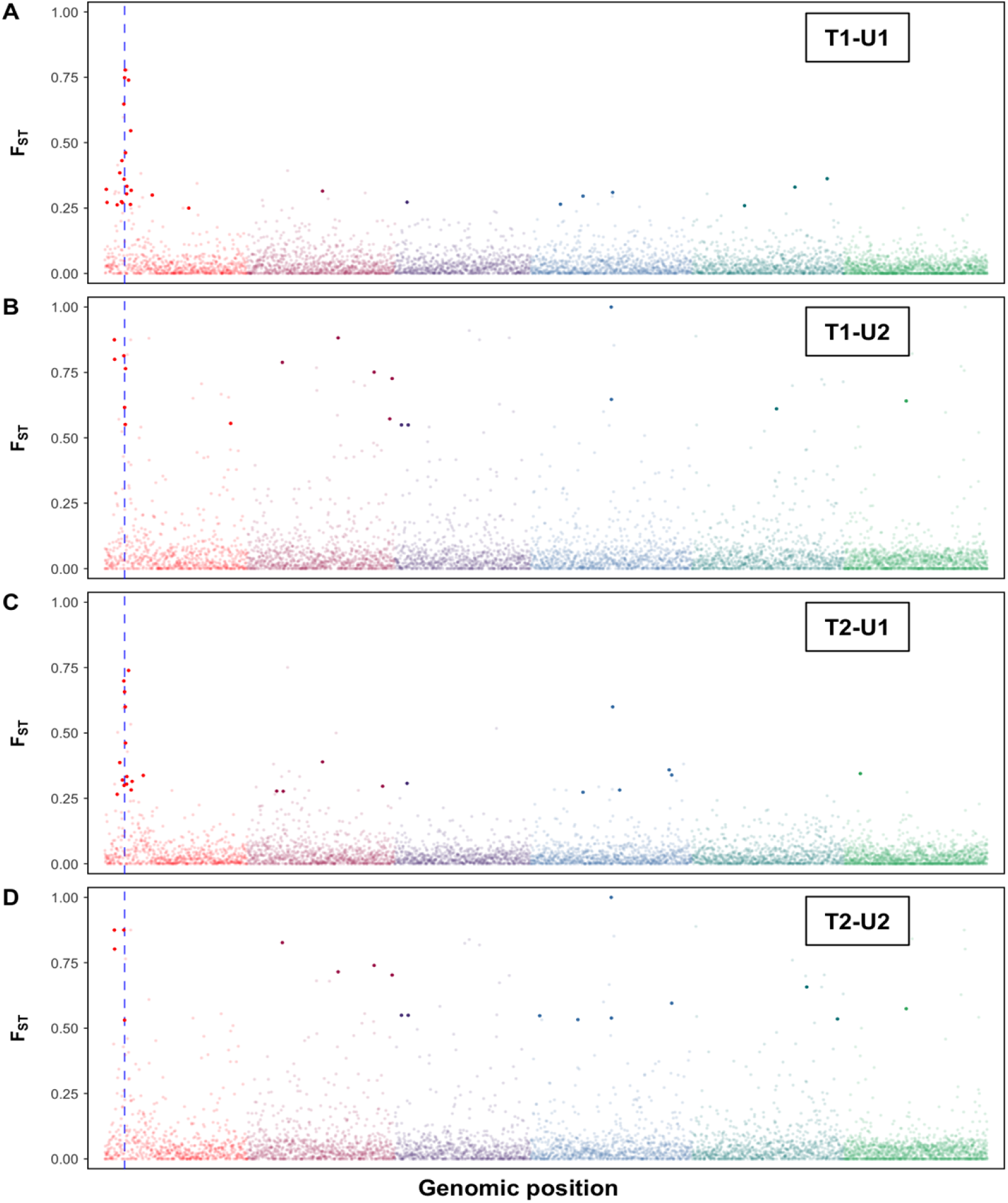
Pairwise F_ST_ analysis between pooled treated and untreated populations. Each dot represents a variable locus between the paired populations, solid colors represent loci with a F_ST_ estimate, as described by the top candidate method [71]. The dashed blue line indicates the position of isotype-1 beta-tubulin.

## 4. DISCUSSION

The small ruminant gastrointestinal parasite *H. contortus* is the leading parasitic nematode model used for anthelmintic resistance research [9,76]. The primary aim of this study was to investigate the number, genomic location(s), and characteristics of the signatures of selection specifically associated with long-term routine use of benzimidazole drugs in *H. contortus* in the field. The widespread use of anthelmintic drugs in livestock, together with high parasite migration as a result of animal movement, makes it difficult to find *H. contortus* populations that have either been subject to selection with a single drug class or have not been exposed to drug selection at all. However, our previous work identified two *H. contortus* populations from government farms in Pakistan (designated T1 and T2 in this paper) that have been subjected to intensive selection pressure by benzimidazoles over many years, but not to selection by other drug classes [52]. In the present work we have compared these with two *H. contortus* populations from rural sheep flocks from the same region (designated U1 and U2 in this paper) that are likely to have had minimal exposure to drug selection [52]. Amplicon sequencing confirmed a high frequency of the previously known F200Y isotype-1 β-tubulin benzimidazole resistance mutation in the two selected populations (T1 and T2) and a very low frequency in the two unselected populations (U1 and U2), consistent with the respective drug selection histories of the populations (**Fig. 1A**). Genome-wide pairwise F_ST_ analysis of 43,587-48,227 RADseq markers confirmed low levels of genetic differentiation between the four populations with U2 being the most differentiated. Consequently, these populations were particularly suitable to investigate the number, genomic location(s) and characteristics of the signatures of selection specifically associated with long-term routine use of benzimidazole drugs in the field. In this paper, we have presented a reduced representation genome-wide approach, in which a large panel of RADseq markers were mapped to the recently completed *H. contortus* chromosomal-scale genome assembly, to identify the main genetic loci under selection by long term use of benzimidazoles these *H. contortus* field populations (Doyle et al., 2020). A single worm genotyping approach was chosen, together with study populations from the same geographical region and minimal genetic differentiation, to maximize sensitivity for detecting signatures of selection in parasite field populations. This experimental design also allowed us to sub-sample in different ways and pool our data in order to investigate how the sensitivity of signal detection varied in terms of sample number and read depth.

### 4.1 The major genomic signature of selection associated with benzimidazole selection in two *H. contortus* field populations surrounds the isotype-1 β-tubulin locus

There is a substantive body of work indicating the importance of a number of non-synonymous mutations in the isotype-1 β-tubulin gene in benzimidazole resistance in *H. contortus* and related ruminant gastrointestinal nematodes of the superfamily trichostrongyloidea [reviewed in 8,77]. This work includes studies that show evidence of selection and high frequencies of codon F167Y, E198A, E198L, and F200Y mutations in benzimidazole resistant parasite populations, and a recent CRISPR/CAS9 reverse genetic study showed that these specific substitutions in the *C. elegans ben-1* β-tubulin gene are sufficient to confer benzimidazole resistance without a fitness cost [78]. There is also evidence to suggest that, although these isotype-1 β-tubulin gene mutations are important determinants of benzimidazole resistance, other loci may also play a role. For example, in *H. contortus,* deletion of isotype-2 β-tubulin gene [31] and increased levels of benzimidazole glycosidation due to increased UDP-glucuronosyltransferase (UGT) expression [79] have been suggested to contribute to resistance. In addition, in *C. elegans,* a QTL on chromosome IV that does not map to the *ben-1* locus, has been shown to underlie the benzimidazole resistance phenotype of some field populations [80], as well as a QTL on the X chromosome [37]. Whole genome-sequencing of adult *H. contortus* worms from different regions of the world confirms historical selection around the isotype-1 β-tubulin locus in addition to many other loci with signatures of selection [38]. Consequently, although isotype-1 β-tubulin gene mutations are clearly important causes of benzimidazole resistance in *H. contortus,* their relative importance with respect to other genetic loci remains a major knowledge gap.

The genome-wide data presented here revealed that the dominant signal of selection for the two different benzimidazole selected *H. contortus* field populations (T1 and T2) was centred on the isotype-1 β-tubulin locus on the left arm of chromosome I. No such signal was present in the two populations with a history of little or no benzimidazole selection (U1 and U2). The selection signal is clearly defined by a region of elevated F_ST_ in pairwise comparisons of the two treated with the two non-treated populations and by reduced expected heterozygosity and elevated linkage disequilibrium in the selected but not the unselected populations. The region of elevated F_ST_ extends across a broad region of the left arm of chromosome I (1,663,419 – 11,549,932) but the regions of reduced nucleotide diversity and elevated linkage disequilibrium are much narrower (π: 5,708,200 – 8,113,930; LD: 6,450,000 – 8,325,000). The isotype-1 β-tubulin gene is located centrally in these regions around position 7,027,095. Overall, these results strongly suggest that the isotype-1 β-tubulin gene is the single most important benzimidazole resistance locus in the *H. contortus* genome of field populations on both government farms examined in this study.

The T1 and T2 *H. contortus* populations may not be completely independent since the flocks were founded by animals from the same region 30 years ago and there may have been some historical animal movement between the farms during the intervening period [52]. Interestingly, the same major isotype-1 β-tubulin resistance haplotypes were detected by short-read amplicon sequencing (**Fig. 1D**). Also, pairwise F_ST_ values of the RADseq markers surrounding the isotype-1 β-tubulin locus were not elevated in pairwise comparisons between the T1 and T2 populations. Together these suggest the same resistance alleles are present in the two populations and so could have common origins. Nevertheless, these two populations have been independently selected by benzimidazole treatment for around 30 years and our data reveals the region surrounding the isotype-1 β-tubulin locus is the most strongly selected in both cases.

### 4.2 Additional genomic regions show signatures of selection after benzimidazole treatments

In addition to the predominant selection signature surrounding the isotype-1 β-tubulin locus in both the T1 and T2 populations, the ddRADseq genome-wide scans detected at least one other region of interest towards the middle of chromosome I (26,568,594 – 26,625,000). This region had a clear signature of selection – as defined by increased population differentiation, reduction in nucleotide diversity and elevated linkage disequilibrium – in the treated but not the untreated populations (**Fig. 2**, **Fig. 3A,B**). This signal is much narrower than that surrounding the isotype-1 β-tubulin locus but is clearly defined by three different measures of selection in both treated populations and not in the two untreated populations. The region encompasses only five genes and may contain a novel drug resistance locus, albeit likely of secondary importance to the isotype-1 β-tubulin gene. Gene ontology revealed no specific functions for the five genes identified in the most narrow region. But a gene (HCON_00018690) with an ABC transporter domain is located close to this region (26,693,160-26,714,246 Mb). Deletion or increased expression of ABC transporters have previously been associated with anthelmintic drug susceptibility or resistance in nematodes, respectively [81,82].

A second region of potential interest is a small region encompassing one gene on chromosome II (**Fig. 2**; region: 13,221,871 – 13,270,135). This region is one bin removed from the bin containing isotype-2 β-tubulin (13,433,296 Mb) and was marked by elevated F_ST_ between both the treated T1 and T2 isolates and U1, but not U2 (**Fig. 2A**). Additionally, this region has reduced nucleotide diversity in T2, but not the other populations (**Fig. 2B**). Short-read amplicon sequencing of a region encompassing codons 167, 198 and 200 of the isotype-2 β-tubulin gene in the T1 and T2 populations did not detect F167Y, E198A, E198L or F200Y candidate resistance mutation in this gene. As a result, the potential role of variation in the isotype-2 β-tubulin gene with regards to resistance in these populations remains unclear.

### 4.3 ddRADseq marker panel development for genome-wide analysis

In this work we explored the utility of a reduced representation genotyping approach, ddRADseq, for detecting signatures of drug selection in a parasitic nematode genome because of its potential cost effectiveness in large scale field studies, even for laboratories with relatively limited budgets. The challenges and opportunities of RADseq to detect loci associated with adaptation have recently been discussed extensively [83–86]. The extent of linkage disequilibrium (LD), the availability of a reference genome, and the resulting marker density needed for detection of adaptation are the main challenges to address. For this study, we generated ddRADseq data from 20 individual worms from each of four populations, following whole genome amplification.

To date, most published applications of ddRADseq on metazoan organisms have involved relatively small marker panels (up to 1 marker per ∼20,900 kb according to a recent review of SNP recovery in aquaculture species [87]) and have successfully been used to study population structure. With regards to genome-wide scans to look for signatures of selection, a major challenge is to develop sufficiently dense marker panels across the genome. For most non-model organisms the extent of LD is unknown, complicating the prediction of the number of markers needed to detect signatures of selection. Another challenge is allelic dropout, which occurs as a result of restriction site heterogeneity, and limits the number of recovered loci that are shared between populations [88,89]. For *H. contortus*, allelic dropout is likely as a result of of its extremely high levels of genetic diversity (reviewed in Gilleard and Redman [90]). We performed an *in silico* digestion with the previously published draft reference genome ([62], GCA_000469685.1) to choose a combination of restriction enzymes and fragment sizes to produce a panel of ∼50k markers that we could recover from at least three of the four populations. At an average of 1 marker per 3kb this represents one of the densest ddRADseq marker sets produced for a metazoan organism to date. Two recent papers have described RADseq SNP marker panels for *H. contortus* with 2,667 [91] and 82,271 [92] markers, respectively. Although the latter study reported a large number of SNPs, this is not comparable to the present study because they reported multiple SNPs per RADseq marker and genotyped pooled genomic DNA to study genetic diversity within and between five populations from different countries.

### 4.4. Considerations to optimize genome-wide approaches to identify signatures of selection in parasitic nematode field populations

Our study and methodology was designed to maximize the sensitivity of detection of the genomic signals of selection associated with benzimidazole treatments of *H. contortus* in the field. The single worm ddRADseq dataset we generated comprised an average sequence depth of >9 reads per marker per sample which we examined in a number of ways to investigate how the sensitivity of detection, based on pairwise F_ST_ analysis and expected heterozygosity, was impacted by the number of individuals, total read depth and reference genome quality. We also investigated the ability of pooled sequence data to detect the signals.

*H. contortus* has substantial population structure between countries and a low but discernible population structure within countries [reviewed by 93]. The identification of genomic signatures of selection in specific regions of the genome can be confounded by population structure [94]. Consequently, the four study populations were taken from the same geographical region to limit the extent of between-population genetic differentiation across the genome. The expected limited differentiation was confirmed by the genome-wide F_ST_ values of <0.05 for all pairwise comparisons (**Table 2**). Further, most of genetic differentiation between the populations was accounted for by chromosome I suggesting it was mainly the result of selection at the isotype-1 β-tubulin locus (**Fig. 1B,C**). Population U2 appeared to be the most genetically distinct of the four populations if the variation on chromosome I is ignored (**Fig. 1C**). There are two alternative explanations for this: (i) genuine genetic differentiation of this population with respect to the others, especially given the relative distance of the sampling location of U2 in Sargodha compared with those of T1, T2 and U1 in Lahore (all four located in the Lahore region), or (ii) a technical artifact as a result of generating the RADseq data for this population on a separate sequencing run as a result of initial poor read recovery of that sample. Fragment size selection during library preparation could have affected the recovered markers. However, based on the markers shared across all populations, this is not a significant concern. Whatever the cause, the strength of the signature of selection around the isotype-1 β-tubulin locus was much lower in pairwise F_ST_ comparisons using the U2 than the U1 population (**Fig. 2**). This illustrates the importance of careful matching of parasite populations being compared and controlling of experimental procedures to minimize genome-wide differentiation and so maximize the sensitivity of detection.

The signature of selection around isotype-1 β-tubulin could be detected with RADseq data from as few as 5 individual worms per population and as little as two reads per marker when 20 individuals were used (**Table 1 and Fig. 4**). Based on the data presented here, aiming for a 5-6.5X coverage of 1.1 to 1.4% of the genome after data cleaning should allow detection of strong signatures of selection. The number of markers expected based on the species of interest and enzyme combination can be calculated *in silico* (Lepais and Weir 2014; Mora-Márquez et al. 2017). For expected heterozygosity, the signatures of selection disappear earlier than for F_ST_; at less than 15 individuals and between 1 and 0.5M reads (**Figs. S7 and S8**). An advantage of expected heterozygosity is that populations do not have to be as carefully matched.

Draft reference genome sequences are becoming available for an increasing number of parasitic nematode genomes (50 Helminth Genomes Project, 50HGP; http://www.sanger.ac.uk/science/collaboration/50hgp). However, there are still very few chromosomal-scale reference genome assemblies for these organisms [but see 66]. Consequently, genome-wide analyses in parasitic nematodes are generally undertaken using draft reference genomes with limited contiguity [95–97]. In order to simulate this situation, we mapped the full set of RADseq read data against a previously published *H. contortus* draft reference genome ([62], GCA_000469685.1). Numerous scaffolds with elevated F_ST_ relative to the rest of the genome were identified as showing a signal of selection (**Fig. S9**). In the absence of positional information for the different genomic scaffolds, this data could be potentially misinterpreted as evidence of multiple signatures of selection being present in the genome when in fact it is due to the hitchhiking effect of a single large selective sweep around the isotype-1 β-tubulin locus. This is an important point to consider when working with draft reference genomes that still have relatively poor contiguity.

We also compared the detection of genomic selection signatures using pooled versus single worm RADseq data. We pooled data from individual samples by merging reads from all samples per population into one sample and selecting enough reads for 100 x coverage. F_ST_ is high around the isotype-1 β-tubulin gene in comparisons with U1, but it is not significant in comparisons with U2 (**Fig. S5**). This result indicates the limited power of F_ST_ estimates in pooled populations if there is some population structure present. For expected heterozygosity no evidence of selective sweeps was detected using the pooled data (**Fig. S6**). The use of pooled data for genome-wide scans can be improved by increasing sample counts [98] but the ability to do this will depend on the number of well-defined isolates and the budget available. Our data provides a framework to balance the relative merits of sequencing a smaller number of worms at low depth from a small number of populations compared with undertaking Poolseq using greater sequencing depth on a greater number of populations.

In summary, this study illustrates the power of genome-wide approaches to identify signatures of selection associated with long term anthelmintic treatment of parasitic nematodes in the field. It has shown the critical importance of minimizing the population structure when selected and unselected populations are being compared and the greater sensitivity achieved by single worm sequencing compared to Poolseq approaches. We have shown that the isotype-1 β-tubulin gene is quantitatively by far the most important benzimidazole resistance locus in the *H. contortus* genome in the two field populations subjected to long term drug treatment. Interestingly however, we have identified two additional loci that likely harbor mutations associated with benzimidazole resistance but are quantitatively of lesser importance than isotype-1 β -tubulin. Further investigation of these regions may reveal new genes associated with benzimidazole resistance.

## ACKNOWLEDGEMENTS

We would like to thank Stephen Doyle, Axel Martinelli, and the members of the HPI training program for their support during various aspects of the project. We would also like to thank WormBase ParasSite for hosting the *H. contortus* genome and annotation. For the purpose of Open Access, the author has applied a CC BY public copyright license to any Author Accepted Manuscript version arising from this submission.

## SUPPLEMENTARY MATERIAL

**S1 File** Sequencing primers

**S2 File** Genes in regions of interest

**S1 Fig.** Marker density across the genome

**S2 Fig.** LD for all populations

**S3 Fig.** Population differentiation with reducing number of individuals; all populations and the whole genome

**S4 Fig.** Population differentiation with reducing read count; all populations and the whole genome

**S5 Fig.** Population differentiation with pooled data

**S6 Fig.** Diversity estimates with pooled data

**S7 Fig.** Diversity estimates with reducing read count

**S8 Fig.** Diversity estimates with reducing number of individuals

**S9 Fig.** All pairwise comparisons with the draft genome

